# Sex chromosome pairing and multivalent associations during meiosis in diploid and polyploid *Silene latifolia*

**DOI:** 10.1101/2025.08.08.669263

**Authors:** Václav Bačovský, Pavla Novotná, Dylan Phillips, Lucie Horáková, Jana Kružlicová, Jana Čížková, Bohuslav Janoušek, Radim Čegan

**Affiliations:** Department of Plant Developmental Genetics, Institute of Biophysics of the Czech Academy of Sciences, Kralovopolska 135, 612 65 Brno, Czech Republic; National Centre for Biomolecular Research, Faculty of Science, Masaryk University, Kamenice 5, 625 00 Brno, Czech Republic; Department of Life Science, Aberystwyth University, Penglais, Aberystwyth, Ceredigion SY23 3DA, UK; Institute of Experimental Botany of the Czech Academy of Sciences, Centre of Plant Structural and Functional Genomics, Slechtitelu 31, 779 00, Olomouc, Czech Republic

**Keywords:** pachytene, Y chromosome, crossover, synaptonemal complex, asynapsis, meiosis, metaphase I

## Abstract

Sex chromosomes undergo various modifications that affect their synapsis during meiosis. While most of the genome achieves full synapsis by the end of pachytene, the non-recombining regions of XY (or ZW) chromosomes often remain asynaptic, and fail to form physical associations at metaphase I. Despite significant progress in animal models, the meiotic behaviour dynamics of plant sex chromosomes remain largely unexplored. In this study, we employed super-resolution microscopy to analyse 3D chromosome organization and the localization of key meiotic proteins. Namely, we studied the dynamics of ASY1, ZYP1, and HEI10, across the leptotene to pachytene stages, and compared sex chromosome behaviour in dioecious *Silene latifolia* with related gynodioecious *S. vulgaris*. Our findings show that both exhibits a class I crossover (CO) frequency comparable to mammals, indicating moderate COs per bivalent and their similar genetic determinants. We document variation in sex chromosome configurations, from rod bivalents in diploids to open-ring tetravalents in autopolyploids, and characterize Y chromosome behaviour across XXY, XXXY, and XXYY karyotypes. These results reveal pronounced variation in pairing and synaptic patterns, even within a shared genetic background. We discuss how these patterns reflect the evolutionary trajectory of the non-recombining region and provide the most detailed cytogenetic analysis of sex chromosome pairing in a plant with evolutionary young sex chromosomes.

## Introduction

Sex chromosomes evolved independently across various taxa in animals and plants (Graves, 2006; Vicoso, 2019; Charlesworth, 2016). Though their independent origin, they exhibit a core set of characteristics that include accumulation of deleterious mutations, recombination suppression, Y degeneration, and evolution of dosage compensatory mechanism (Wright *et al*., 2016; Muyle *et al*., 2021; Kratochvíl *et al*., 2021). Following successive steps of recombination suppression, the sequence divergence allows to form “evolutionary strata” (stratum= regions that stopped recombining at different times, and their age is inferred from the level of synonymous substitution between X- and Y-linked allele) (Olito and Abbott, 2025). Deleterious mutations disrupt the reading frame of the Y-linked genes (Bergero and Charlesworth, 2009) and, together with the accumulation of transposable elements (TEs), cause a decrease in the expression of other mutated sequences, leaving the heterologous chromosome weak (Lenormand *et al*., 2020; Muyle *et al*., 2022). In this evolutionary stage the non-recombining region often occupies a significantly large part of the Y (W), which in time, becomes heterochromatic and epigenetically inactive (Sardell and Kirkpatrick, 2019; Wright *et al*., 2016).

The genetic and morphological characteristics of sex chromosomes have significant implications for their meiotic behaviour (reviewed in Page *et al*., 2006). Meiosis, a process in which sexually reproducing organisms form gametes, in plants spores (sperm and egg cells), that contain half the chromosome number and, display canonical features across different phylogenetic groups (Harrison *et al*., 2010). The variability among subsequent generations is produced by recombination during prophase I which involves several key steps that also ensure faithful chromosome segregation. This includes chromosome synapsis via the formation of a protein scaffold along the chromosomes (SC), the formation of double strand breaks (DSBs) and DNA exchange (crossovers, COs), and the separation of homologous chromosome pairs (bivalents)(Mercier *et al*., 2015). Most notably, the process of synapsis is completed at the end of zygotene of prophase I. The proper meiotic pairing, manifested as full synapsis, is typically associated with DNA exchange. COs are later resolved at the diplotene-diakinesis stage (Qiao *et al*., 2012; Mercier *et al*., 2015). To prevent the transmission of damaged or improperly organized genetic material found on sex chromosomes, asynapsed (unsynapsed) chromosomes are subjected to the silencing mechanisms termed as meiotic silencing of unsynapsed chromatin (MSUC) (Turner *et al*., 2005). This process ensures the chromosomes that fail to synapse are transcriptionally silenced, preventing the expression of potentially harmful genes that might be present on unpaired chromosomes and thereby reducing the risk of gametogenic failure (Cloutier and Turner, 2010). In eutherian mammals, asynapsis is a prominent feature of non-homologous regions of the X and Y chromosomes (Turner, 2007; Page *et al*., 2006). The asynapsed regions are subjected to the process of meiotic sex chromosome inactivation (MSCI) and it may involve changes to the tripartite proteinaceous axis of the synaptonemal complex (SC), particularly the axial and central elements, which may undergo structural and functional modifications (Ishiguro, 2023; Cahoon and Libuda, 2019). These architectural alterations contribute to sex-specific differences in recombination landscapes and may have profound consequences for meiotic stability, CO formation, or in some cases affect fertility (Sardell and Kirkpatrick, 2019; Alavattam *et al*., 2021). While such phenomena have been extensively studied in animals, the behaviour of sex chromosomes in plants remains largely underexplored. In fact the asynaptic behaviour of sex chromosomes, to date, have only been described only in the genus *Rumex*, namely *R. suffruticosus, R. acetosella* and *R. acetosa*, which display distinct XY or XYY meiotic configurations (Cuñado *et al*., 2007). Interestingly, *Vasconcellea parviflora* represent an early evolutionary stage of sex chromosomes and appears to exhibit limited asynaptic behaviour in the heterochromatic domain on the Y chromosome (Iovene *et al*., 2015). However, the dynamics during prophase I remain unknown.

Dioecious plants offer a unique opportunity to study meiotic synapsis because the sex chromosomes are still ongoing degenerative processes and are of independent origin (Garcia *et al*., 2023). The dioecious plant model, *Silene latifolia*, possesses a large heteromorphic sex chromosomes that evolved 11 MYA (Moraga *et al*., 2025). Both sex chromosomes have accumulated a large proportion of tandem repeats, some of which are X- or Y-chromosome specific, and some that are predominantly enriched in gene regulatory regions (Akagi *et al*., 2025; Moraga *et al*., 2025). The sex chromosomes display modest heterochiasmy, between linkage groups (LGs) 5, 6, and 11 with higher recombination in males compared to females (Filatov, 2023). The recombinationally inactive regions are highly enriched with LTR-Gypsy and LTR-unknown class of transposable elements, which is similar to the observed phenomenon in related species with no sex chromosomes, *S. vulgaris* (Akagi *et al*., 2025). Despite the large developments in the genomic resources available for this model plant, the knowledge of Y chromosome pairing behaviour, sex chromosomes association during prophase I and progression through meiotic division remain limited.

To uncover the sex chromosome dynamics of *S. latifolia*, we used high resolution microscopy and 3D specimen preparation technique to study XY pairing during prophase I. We developed a polyclonal antibody to label the ZYP1 proteins that forms the transverse filaments of the SC of *Silene*, and visualized ongoing synapsis between autosomes and sex chromosomes. Next, we generated artificial triploids and autotetraploids in *S. latifolia* and analysed the XY pairing in the various polyploid backgrounds. We specifically aim to answer (i) whether the sex chromosome dose impairs sex chromosome pairing and synapsis, and (ii) what is the behaviour of the Y chromosome in the presence of another homologous partner. Using a XY-oligo painting probe, we show preferred associations of various sex chromosome doses during metaphase I and discuss the Y chromosome pairing capability that would have implications for the evolution of non-recombining region.

## Methods

### Plant material

Seedlings of *S. latifolia* inbred population U16, made by 16 generations of full-sib mating, were used as a parental population, as described previously in Bačovský *et al*. (2020). The ecotype of *S. vulgaris* originating from Brno region was used as an outgroup. The seeds of both species are owned by the Institute of Biophysics of the Czech Academy of Sciences. Plants were kept in a greenhouse under a 16 h light/8 h dark cycle at 24°C in controlled conditions.

### Polyploid generation

Triploid and tetraploid individuals were generated using a previously published protocol (Jones *et al*., 2008), using 5 µM oryzalin and 5 mM colchicine. Due to the low survival rate of young, treated seedlings (Supplementary Note S1, Table S1), we used the following protocol treating female flowers 2 days after pollination (DAP) (Table S1). Diploid female flowers of U16 generation were first fertilized with diploid U16 male flowers, and 2 DAP the same flowers were washed for 1 h in deionized water, incubated for 7 h in 5 mM colchicine solution, and washed again in deionized water for 1 h. After 14+ days, mature seeds from the treated flowers were sterilized as described previously (Bačovský, Čegan, *et al*., 2022). Seeds were grown in a greenhouse under the same conditions as the parental plants. Mean DNA nuclear content of adult individuals was analysed with flow cytometry, and fluorescence *in situ* hybridisation (FISH) using a previously described protocols (Bačovský *et al*., 2020; Bačovský, Janíček, *et al*., 2022). For determination of the number of sex chromosomes within the polyploid individuals, we used two satellites – subtelomeric (X43.1) and centromeric (STAR-C) satellites, and X-chromosome pseudoautosomal region (PAR) oligo painting probe (XY-PAR) as described in Bačovský *et al*. (2020). Hermaphrodite flowers of tetraploid individuals were self-crossed and were used to generate the F2 population. Subsequently, the same plants were used as pollen donors and crossed with U16 diploid females to produce triploid individuals.

### Genome size measurement

The nuclear genome size of tetraploid and triploid individuals were estimated following the protocol described in Doležel *et al*. (2007). Briefly, fresh leaf tissue of *S. latifolia* or *S. vulgaris* and the internal reference standard were chopped together in a 1 mL volume of Galbraith’s buffer using a razor blade (Galbraith *et al*., 1983). The crude suspension was filtered through a 50 µm nylon mesh. The filtrate was made up to 50 µg/mL propidium iodide and 50 µg/mL RNase. Samples were analysed using a CytoFLEX flow cytometer (Beckman Coulter, United States) equipped with a 488 nm blue laser. *Pisum sativum* ‘Ctirad’ (2C = 9.09 pg; Doležel *et al*., 1998) and *Solanum lycopersicum* ‘Stupicke polni tyckove rane’ (2C = 1.96 pg; Doležel *et al*., 1992) were used as internal standards. Each sample was analysed three times, each time on a different day. 5000 nuclei per sample were analysed and 2C DNA contents (in pg) were calculated from the means of the G1 peak positions by applying the formula: 2C nuclear DNA content = (sample G1 peak mean) × (standard 2C DNA content)/(standard G1 peak mean). Mean nuclear DNA content (2C) was then calculated for each accession. DNA contents in pg were converted to genome lengths in bp using the factor suggested by (Doležel *et al*., 2003), i.e., 1 pg DNA = 0.978 Gbp (Table S1, Fig. S1).

### ZYP1 antibody development

*Hordeum vulgare* and *Arabidopsis thaliana* sequence of ZYP1 gene sequences (Higgins *et al*., 2005) were used to search *Silene* orthologs in NCBI database (Fig. S2a). Based on the quality and sequence length, we designed primers for *S. colpophyla* ZYP1 sequence, annotated in Geneious Prime (2023.1.1). The whole sequence was amplified from *S. latifolia* gDNA using SlZYP1 F1/R1 primers, (SlZYP1 F1 – TGGCTAGGTCTCGAGTCGAA and R1 – CTCAGCTTGGCGATTGATGC), synthesized by GeneriBiotech (Hradec Kralove, Czech Republic). The gene was amplified using Q5 High-fidelity DNA polymerase (M0491S; NEB), following the manufacturer instructions. Resulting PCR products were purified using QIAquick PCR Purification Kit (28104, QIAGEN), and used for blunt cloning using CloneJET PCR Cloning Kit (K1231; ThermoFisher). Ligation reactions were incubated overnight at 16 °C. After desalting, competent *E. coli* cells were transformed by electroporation using a MicroPulser Electroporator (Bio-Rad). Transformed cells were allowed to recover for 30 minutes and then plated on LB agar supplemented with ampicillin (100 mg/L). Plates were incubated overnight at 37 °C. The following day, PCR on resulted colonies was performed using pJET1.2-specific primers to identify positive clones. Individual bacterial colonies were used as templates. Primers and nucleotides were removed using ExoSAP reaction. Sequencing primers (pJET1.2 F/R) were used in all reactions. Samples were sequenced by Macrogen (Amsterdam, Niederlands) and aligned in Geneious Prime (Table S2, Fig. S2b). The whole consensus protein sequence was analysed by Kyte & Doolittle algorithm with linear weight variation model (Kyte and Doolittle, 1982), and compared to barley and Arabidopsis (Fig. S2c, Table S2). To verify the protein structure and similarity to published ZYP1 sequences, the whole protein sequence was computed by AlphaFold2 (Jumper *et al*., 2021) (Fig. S2d, e). Following the verification steps, the whole protein was used for immunization of two Guinea pigs to generate a řřmt complex polyclonal antibody. The codon optimization, protein synthesis, immunization, and antibody purification were performed by GenScript (New Jersey, USA).

### Chromosome preparation and meiotic analysis

*S. vulgaris* (2n = 24), *S. latifolia* U16 (2n = 2x = 24, XY), triploid U17 (2n = 3x = 36, XXY) and autotetraploid U17 (2n = 4x = 48, XXYY; 2n = 4x = 48, XXXY) were used for further meiotic analysis (Tables S1). Chromosomes in metaphase I were prepared from young flower buds. Anthers were dissected under Olympus Stereomicroscope SZX16 (Evident) supplemented by LED fluorescence and light system, and Olympus DFLPLAPO 0.8PF objective. Anthers were first squashed in 1% acetocarmine and desired stage was verified under Light Microscope CX43 (Evident), supplemented with PlanC 40x objective (NA 0.65) and phase contrast. The remaining anthers of the same size within the same flower were placed into Clarke’s fixative (ethanol: glacial acetic acid, 3:1, v:v) for 72 h at RT, washed with freshly prepared Clarke’s fixative again, and stored at −20°C until use. Chromosomes for FISH experiments were prepared by the squashing technique as described in Bačovský, Janíček, *et al*. (2022), with minor modification. Briefly, the incubation time in the enzymatic mixture was 20 min and slides with a well-preserved number of chromosomes were stored in 96% EthOH at −20°C until use. The FISH was performed using the same combination of probes as for karyotype screening: X43.1, STAR-C, and XY-PAR oligo painting probes. Chromosome pictures were captured with Olympus AX70 microscope equipped with CCD1 camera, fluorescence PhotoFluor LM-75 light source, PlantAPO oil objective 100x (NA 1.35) and filter set for 405, 488, 561 and 642 nm fluorescent wavelength. All channels were processed in Adobe Photoshop v. 26.9.0.

### Immunostaining and super-resolution microscopy

Anthers in prophase I were screened in the same way as for metaphase I preparation. Specimens for 3D immunolocalization were prepared as previously described in Hurel et al. (2018), with minor modifications. Briefly, anthers from young flower buds were fixed in 2% (w/v) paraformaldehyde, macerated with a brass rod in buffer A and embedded in acrylamide. Embedded meiocytes were washed in washing buffer (1xPBS supplemented with 1% Triton X-100 and 0.5mM EDTA) for 2 h, and incubated with following polyclonal antibodies diluted in 1% blocking buffer (1% BSA (w/v) in 1xPBS, 0.1% Tween20): α-ASY1 (rat, 1:200; Hurel et al., 2018), α-HEI10 (1:200; (Chelysheva *et al*., 2012), α-SlZYP1 (guinea pig, 1:300; this study). After incubation with primary antibodies (48h) at 4°C, embedded meiocytes were washed 4x 1h in 1xPBS at RT and incubated with secondary antibodies conjugated with Fluorescein (donkey α-rat; 712-097-003), Cy3 (goat α-rabbit; 111-167-003) or Alexa 647 (donkey α-mouse; 715-606-003), diluted 1:400. Immunostained cells were imaged using structured illumination microscopy (3D-SIM^2^) on a Zeiss Elyra 7 microscope equipped with metal halide HXP 120 V fluorescence light source, Plan-apochromat 63x oil objective (1.46 NA), filter set for 405, 488, 561 and 642 nm fluorescent wavelength and 2xPCO edge sCMOS camera with pixel size resolution 6.5 µm x 6.5µm at CELLIM facility (Brno). Images were captured in ZEN Black software with SIM^2^ deconvolution.

### Image analysis and HEI10 classification

For image deconvolution in ZEN Black, we applied adjusted parameters with standard-fixed values for ASY1 and ZYP1 across all three meiotic stages, using the scale to raw image option in advanced histogram settings. For improved detection of HEI10 signals during leptotene and zygotene, the regularization weight was set to 0.03. In pachytene, HEI10 signal was processed using strong-fixed values and again scaled to raw image histogram settings.

Subsequent image analysis was performed using Imaris (v10.0, Oxford Instruments). To segment the synaptonemal complexes (SCs), surface creation was applied using a smooth surface detail value of 0.05 and absolute intensity thresholding. The filter surface tool was manually adjusted (semi-automatic mode) to retain all relevant ASY1 and ZYP1 signal intensities, with no additional subclassification of detected objects. Leptotene, zygotene, and pachytene stages were identified based on ASY1/ZYP1 distribution and the number of HEI10 foci.

HEI10 foci were classified according to their size and signal intensity as described by Randall *et al*. (2022), with minor modification and differentiating only two types. We used the shortest distance algorithm for spot detection, with either variable spot sizes or a fixed estimated x, y-diameter of 0.150 µm. Point spread function (PSF) elongation along the z-axis was modelled with a diameter of 0.350 µm. These stricter size estimates, compared to Randall *et al*.(2022) were calculated by measuring HEI10 foci in x-y-z projections from slice view mode in ten representative nuclei. Foci were further filtered by mean intensity and then classified into type A or type B based on both maximum intensity and average x/y-diameter: type A foci were <0.07 µm, and type B foci >0.08 µm. Once parameters were optimized for each meiotic stage, all images were processed using the batch analysis tool. Quantitative data were exported to Excel and further analysed in R Studio. Statistical significance of class I crossover differences was assessed using two-way ANOVA followed by Tukey’s multiple comparisons test.

## Results

### Sex chromosome configurations vary with ploidy level and XY chromosome dose

To understand how sex chromosome composition and ploidy level influence meiotic chromosome pairing in *Silene*, we examined male meiocytes at metaphase I across diploid (*S. vulgaris, S. latifolia*), and (derived) artificial triploid (XXY), and autotetraploid (XXXY and XXYY) individuals. Plants with higher ploidy level were derived from flower buds treated with 5mM colchicine 2 DAP (Fig. S1, Table S1, Note S1). We systematically reconstructed the autosomes configurations and sex chromosome associations during metaphase I by 2D specimen preparation with previously published oligo-painting probe (XY-PAR), and subtelomeric (X43.1), and centromeric (STAR-C) repeats. This combination of probes allows to differentiate three orthologous autosomes in related *S. vulgaris* (2n = 2x = 24), and to position the pseudoautosomal region (PAR) on the q-arm of Y chromosome and on the p-arm of X chromosome in *S. latifolia* (2n = 2x = 24, XY) (Fig. 1). This XY-PAR positioning was consistent with a previously published models for both species (Bačovský *et al*., 2020). Cytogenetic observation of the probes in the neotriploid and neotetraploid *S. latifolia* confirmed their order was preserved (Fig. S3–S6). In *Silene vulgaris*, the three orthologous autosomes identified by XY-oligo painting probe consistently formed two ring bivalents and one rod bivalent (Fig. 1, S3). In contrast, male plant of *S. latifolia* formed rod bivalent between the X–Y chromosomes, with the physical association and CO confined to the PAR (Fig. 1, Fig. 2, Fig. S3). Additionally, the subtelomeric repeat (X43.1) was positioned within three other autosomal bivalents in *S. vulgaris* (Fig. 1, S3), and not colocalized with the XY-PAR oligo painting probe.

**Figure 1.**
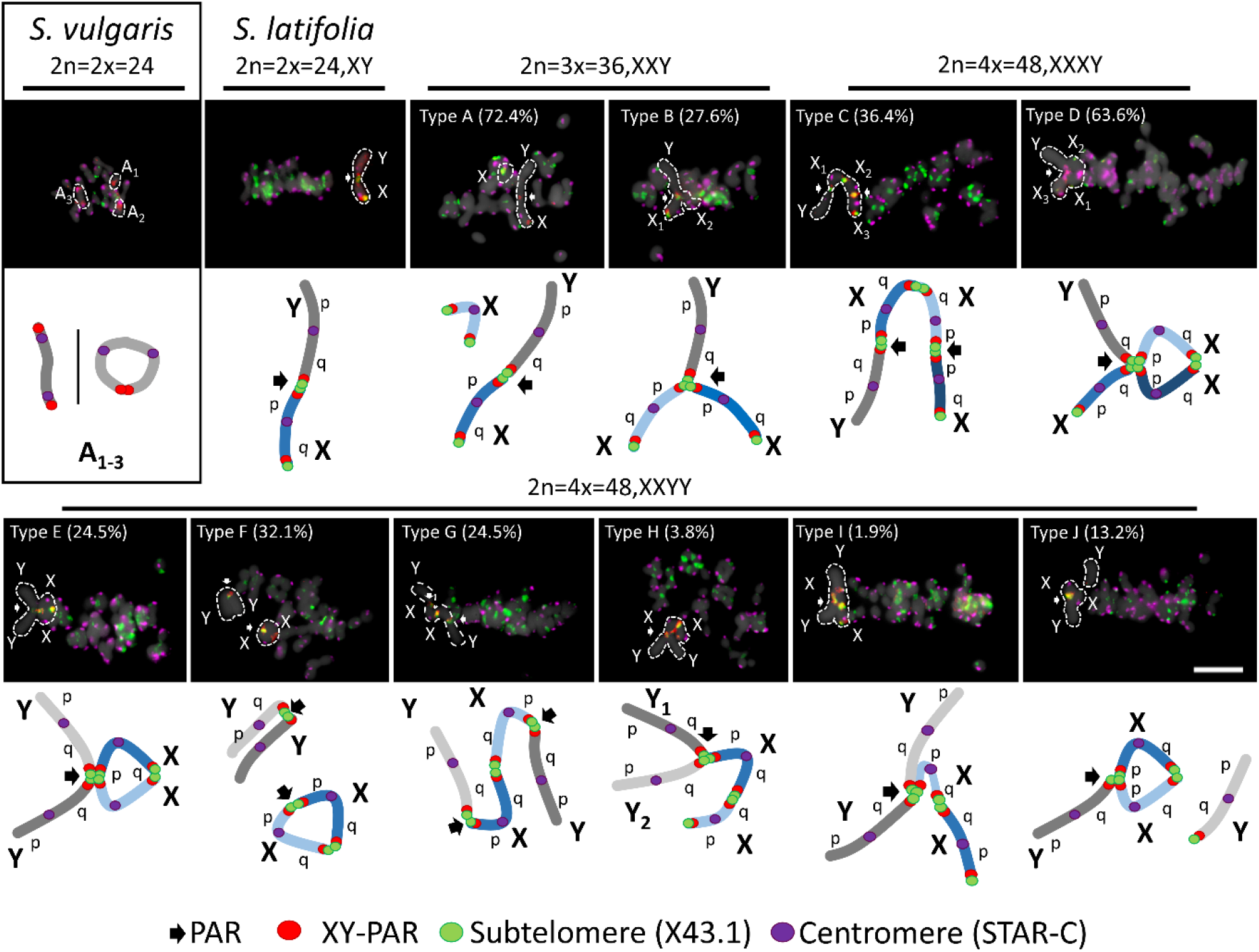
Meiotic segregation and sex chromosome pairing during male metaphase I in *Silene*. In *S. vulgaris*, three autosomes differentiated by XY-PAR positioning, form two ring bivalents and rod bivalent configurations at the metaphase I plate. The three regions are orthologous to XY chromosomes (red). In contrast, in *S. latifolia*, the X and Y chromosomes consistently form a rod bivalent at the metaphase I plate. In cases of multiple sex chromosomes (2n = 2x = 36, XXY; 2n = 4x = 48, XXXY/XXYY), the sex chromosomes form from left to the right (the upper line): type A – a single rod bivalent and physically separated univalent (X-Y and X), type B – a Y-shaped trivalent (X-X-Y), type C – a chain-like (Y-X-X-X), and type D – an open-ring tetravalent with three Xs and single Y. From left to the right (bottom line): type E – an open-ring tetravalent, type F – a single bivalent (X-X) and rod bivalent structure (Y-Y), type G – a chain-like tetravalent (Y-X-X-Y), type H – an open-ring tetravalent in which two X chromosomes are connected through the q-arm (Y-Y-X-X), or type I – all four sex chromosomes connected via the PAR (YY-X-X). Type J consists of a chain tetravalent structure with a loosely associated Y chromosome (Y-X-X/Y). The formation of all displayed configurations was consistently observed in individual analysed cells (at least 20 per slide) across eight replicates. Sex chromosomes in *S. latifolia* and polyploid individuals were identified using a combination of subtelomeric (X43.1; green) and centromeric tandem repeat (STAR-C; magenta) markers. Arrows indicate PAR. Chromosomes were counterstained with DAPI (gray). Scale bar = 10 µm.

**Figure 2.**
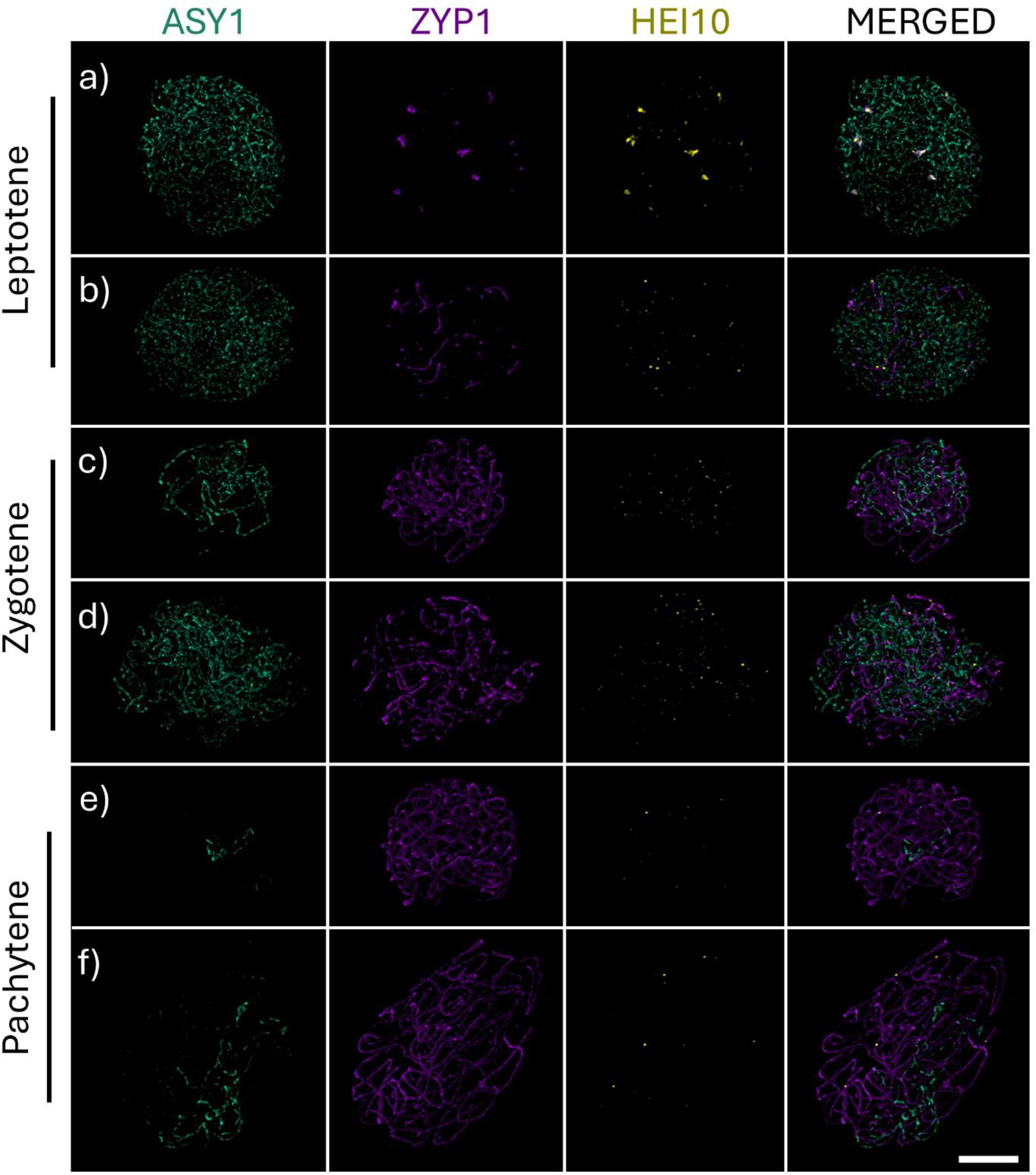
Formation of synaptonemal complexes and immunolocalization of meiotic proteins ASY1, ZYP1, and HEI10. (a–f) The progression of prophase I in *S. vulgaris* (a, c, f) and *S. latifolia* (b, d, f) is visualized through the dynamic localization of ZYP1 along the axial elements. (a, b) During leptotene ASY1 (cyan) labels the axial elements, and central elements (ZYP1, magenta) become increasingly distinct. HEI10 forms numerous small foci along the chromosomes. (c, d) In zygotene, the ASY1 signal becomes fragmented as synapsis progresses, marked by the extension of ZYP1 along the central element, facilitating synaptonemal complex (SC) assembly. HEI10 foci subsequently coalesce into large foci. (e, f) By pachytene, ASY1 (axial elements) was restricted to asynaptic regions of the nucleus, while fully assembled SCs were observed between homologous chromosome pairs. These are three autosome bivalents in *S. vulgaris*, and PAR in *S. latifolia*. HEI10 foci become fewer and larger, indicating a process of coarsening, marking future crossover events. The behaviour of ZYP1, ASY1, and HEI10 was consistent across all analysed cells (n = 74; Table S3). Scale bar = 4 µm.

In triploid *S. latifolia* males (2n = 3x = 36, XXY), two configurations were observed at metaphase I: the predominant type A configuration, where one X chromosome remains univalent while the other pairs with the Y, forming a rod bivalent (72.4%). In the second type B configuration, all three sex chromosomes link at the PAR forming a trivalent Y-shaped association (27.6%) (Fig. 1, S4).

Autotetraploid individuals (2n = 4x = 48) with XXXY or XXYY karyotypes showed a wider diversity of metaphase I configurations (Fig. 1, S5, S6). In XXXY plants, chain-like (type C) and open-ring tetravalents (type D) configurations were frequently observed, indicating at least 2 to 3 CO points among the X chromosomes and their association with the Y (Fig. S5). In XXYY individuals, at least five distinct configurations were detected (types E–J), ranging from open rings and chains to bivalent combinations, often with asymmetrical Y chromosome involvement (Fig. S6).

Among the observed configurations, the open-ring tetravalent was one of the most recurrent and structurally diverse formations. In XXXY individuals, the open-ring tetravalent (type D) was the most frequent configuration, accounting for 63.6% of observed cells (Fig. 1, type D). This configuration typically involved a ring bivalent of two Xs connected at both arms and a third X linked to the Y via the PAR, linked together with the Xs bivalents. Thus, this configuration of XXXY resulted in an open-ring tetravalent (Fig. 1, S5).

In XXYY individuals, open-rig tetravalents appeared in multiple structural variants, including types E and F, which together represented over 50% of observed cells (Fig. 1, S6). These included configurations where two X chromosomes were connected via the q-arm, and one or both Y chromosomes were associated with Xs via the PAR, located on the X p-arm. Additionally, chain tetravalents, types G and I, formed associations in which Xs pair via q-arm, and both Ys were paired via PAR at X p-arm. Overall, the recurrent presence of open-ring tetravalents across different polyploid karyotypes with 3 Xs or 2 Xs suggested that this structure may represent a preferred mode of synapsis and CO resolution under conditions of increased sex chromosome dose and partial homology. In this regard, type H and type I were the least preferred pairings as they were observed in 3.8% and 1.9% of cells, respectively. All configurations were reproducible across biological replicates (n ≥ 20 cells per slide; 8 replicates), demonstrating stable yet diverse meiotic behaviour in response to increasing sex chromosome complexity.

### Progression of synapsis from leptotene to pachytene in relation to sex chromosome dose

To further investigate the configurations observed at metaphase I in diploid, triploid, and tetraploid *S. latifolia* (Fig. 1), we examined the dynamics of meiotic pairing during early prophase I–specifically at the leptotene, zygotene, and pachytene stages (Fig. 2a–f). In both diploid *S. vulgaris* and *S. latifolia*, loading of ASY1, a marker associated with axial element formation, occurred along all autosomes and sex chromosomes (Fig. 2a, b), accompanied by the dot-like emergence of central element signals (ZYP1). As meiosis progresses in zygotene, ZYP1 extends along axial elements, and numerous HEI10 foci become visible, gradually increasing in size from zygotene into pachytene (Fig. 2c–f). By pachytene, most chromosomes were fully synapsed, as indicated by continuous ZYP1 signals and gradual disappearance of ASY1 (Fig. 2e, f).

Three-dimensional (3D) reconstructions of pachytene nuclei revealed persistent asynaptic regions: six distinct ASY1-positive axes corresponding to three autosomal bivalents in *S. vulgaris*, and a single unpaired region composed of two ASY1-marked axes in *S. latifolia* corresponding to the X and Y chromosome (Fig. 2, Fig. 3a, b). These unpaired regions are positioned adjacent to a short, fully synapsed domain marked by ZYP1, corresponding to the PAR. Dissection of individual Z-stacks confirmed the presence of discrete ASY1-ZYP1 transition zones in all three orthologous chromosome pairs (Fig. 3c–e), as well as a well-defined PAR boundary (PAB) in *S. latifolia* (Fig. 3f).

**Figure 3.**
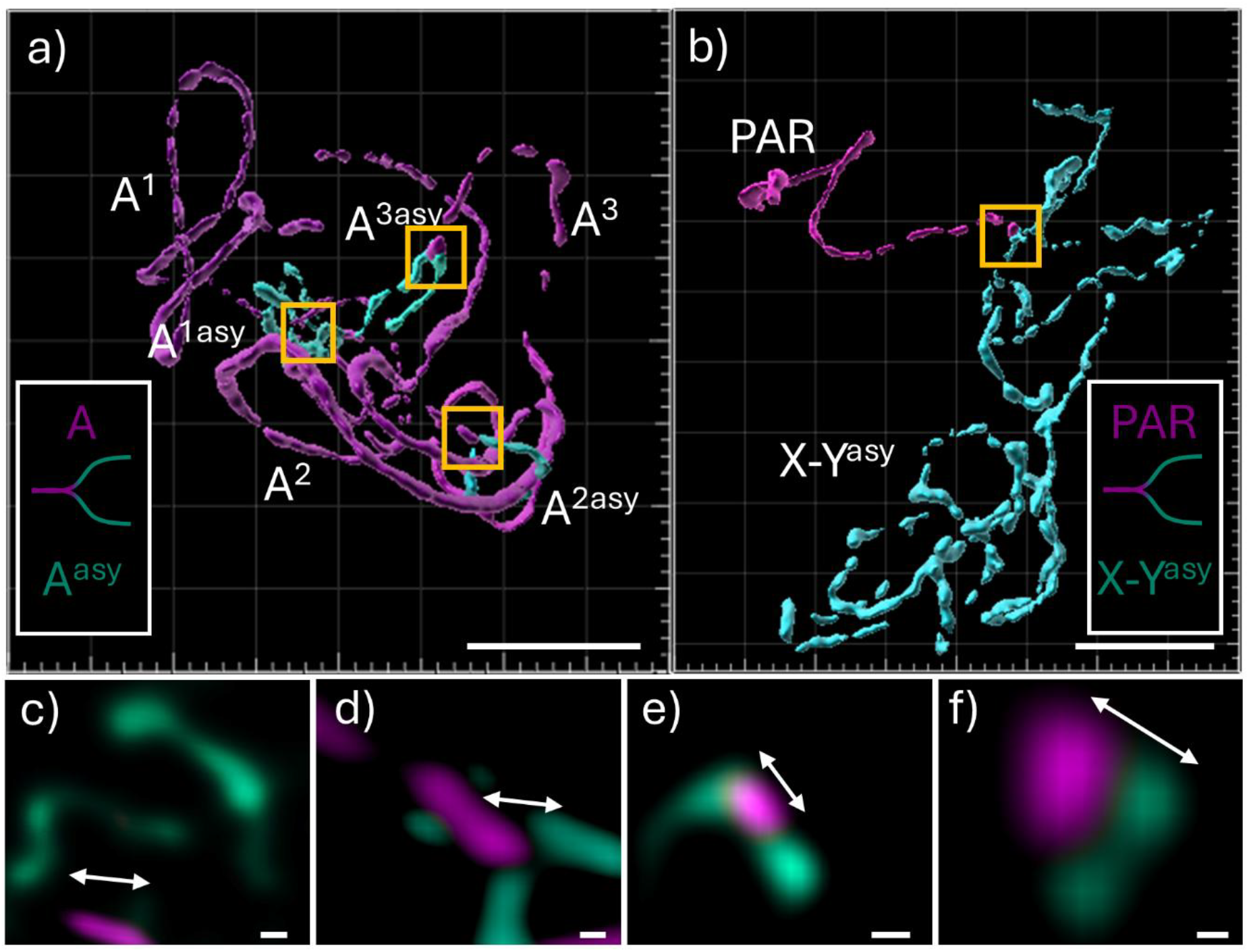
3D analysis of meiotic pairing in pachytene chromosomes of *S. vulgaris* and *S. latifolia*. (a) In *S. vulgaris*, three orthologous chromosome pairs, depicted by ZYP1 (magenta) axis (A^1^ – A^3^), links to three asynaptic regions (cyan) designated as A^1asy^, A^2asy^, and A^3asy^. (b) In *S. latifolia*, a single large asynaptic region of two axis (X–Y^asy^) corresponds to the X and Y chromosomes. PAR is visualized by ZYP1 immunolocalization. White boxes in (a, b) represents the boundary of asynaptic region in both species. (c, d, e) The insets (yellow boxes) of asynaptic regions in *S. vulgaris* (a), the boundaries between synapsed and asynaptic regions are sharply defined, as revealed in maximum intensity projections, marked by arrows. (f) The inset (yellow box) of asynaptic region in *S. latifolia* (b), the asynaptic region displays a clear boundary at the PAR (PAB), extending into two protruding ASY1-marked axes representing the unsynapsed portions of the X and Y chromosomes. Scale bars in main panels (a, b) = 3 µm. Scale bar in insets (c–f) = 0.5 µm.

The same synaptic dynamics were observed in polyploid individuals, including XXY, XXXY, and XXYY genotypes, where clearly distinguishable asynaptic regions persisted despite increased sex chromosome dose (Fig. 4, Figs S7 and S8). The use of 3D imaging nucleus segmentation, followed by axis simulations, allowed us to quantify the ASY1/ZYP1 ratio per cell across different ploidy levels (Fig. 5). As expected, the volume of ASY1/ZYP1 signal increased with genome size, accompanied by a proportional increase in HEI10 foci per cell (Fig. S9). In line with the immunolocalization results, volumetric quantification of the ASY1/ZYP1 differences across leptotene–zygotene–pachytene stages confirmed progressive SC assembly. However, the relative size of the asynaptic region, measured by the ASY1 signal, remained unchanged regardless of increasing sex chromosome dose, and the presence of another X or Y homologous partner in XXXY/XXYY individuals (Fig. 5a, b). This observation aligns with the dominant presence of open-ring sex chromosome configurations at metaphase I (Fig. 1), typically involving partially synapsed X chromosomes with Y chromosomes attached solely via the PAR. The same holds true in the case of XXYY plants in which two Ys are attached in the PAR to the Xs bivalent, forming open-ring or chain-like configuration (type E, Fig. 1). It should be stressed that the chromosome configuration as showed in Fig. 1 could not be differentiated at pachytene and correlated to the ASY1/ZYP1 measurements taken from the 3D specimens.

**Figure 4.**
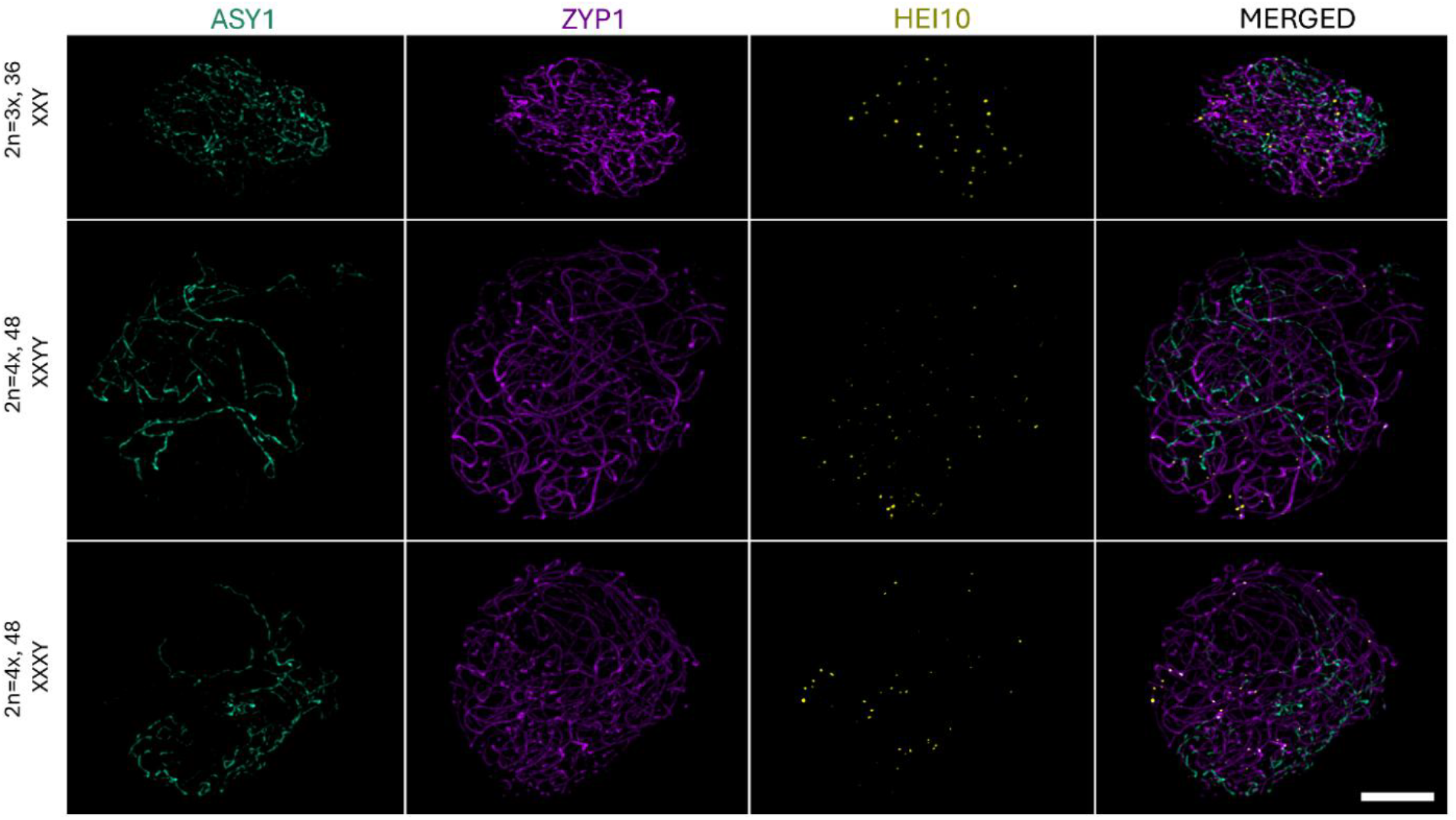
Synaptonemal complex formation in male triploid and autotetraploid *Silene latifolia* individuals. During pachytene, in triploid and autotetraploid plants with XXY, XXYY, and XXXY sex chromosome constitutions, ASY1 (cyan) is confined to asynaptic regions of the nucleus, while fully assembled SCs are present between homologous chromosome pairs (ZYP1; magenta). HEI10 (yellow) foci are observed exclusively on fully assembled SCs. Scale bar = 4 µm.

**Figure 5.**
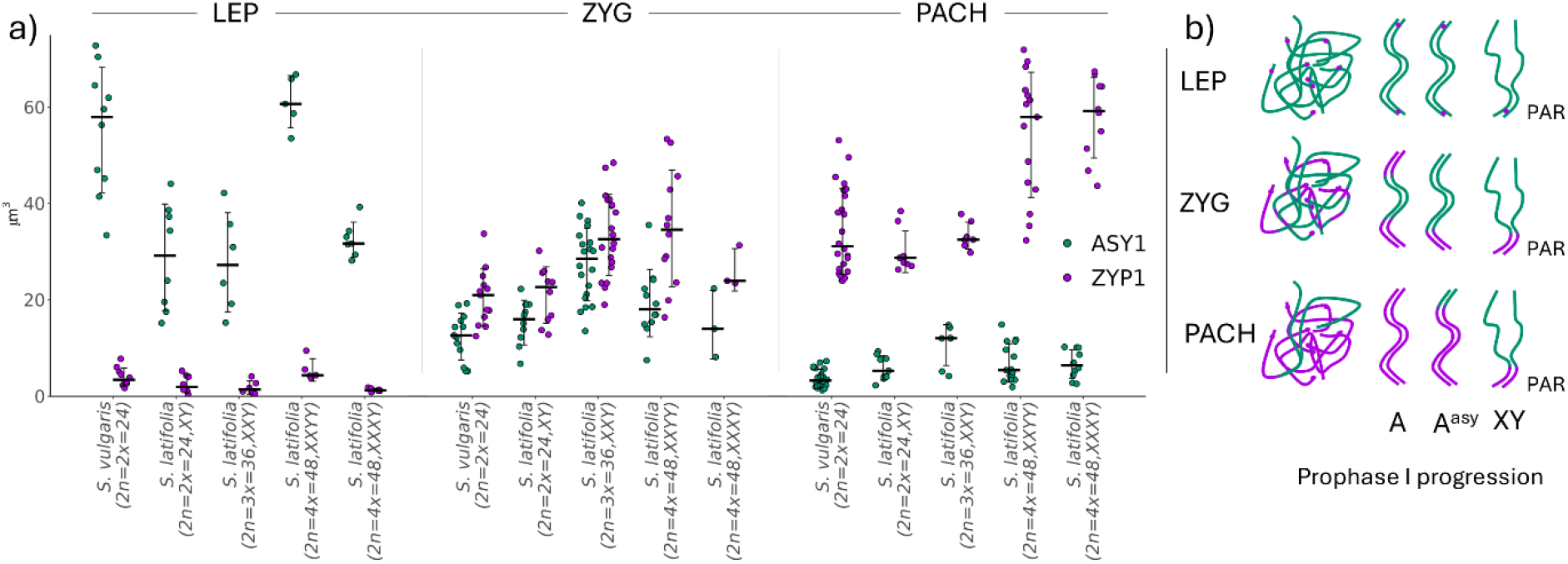
Dynamics of ASY1 and ZYP1 loading during meiotic pairing in prophase I. (a) Quantification of ASY1 (cyan) and ZYP1 (magenta) signal volumes during leptotene (LEP), zygotene (ZYG), and pachytene (PACH) in *S. vulgaris, S. latifolia*, and polyploid individuals. A progressive increase in ZYP1 signal corresponds with a reduction in ASY1 as synapsis proceeds. Despite differences in ploidy and sex chromosome constitution, the overall ASY1-to-ZYP1 transition pattern is conserved. (b) Schematic representation of ASY1 and ZYP1 loading during leptotene-to-pachytene progression in prophase I. In *S. vulgaris*, three autosomal pairs (A^asy^) display asynaptic behaviour. In *S. latifolia*, as well as in XXY, XXXY, and XXYY individuals, the non-homologous regions of the X and Y chromosomes (X–Y^asy^) are asynaptic, marked by strong ASY1 adjacent to short ZYP1-positive PAR segments.

### Quantification of HEI10 foci during meiotic prophase I reveals temporal and karyotype-specific dynamics

The HEI10 foci were classified from leptotene to pachytene into two types – type A unmature HEI10 foci (<0.07 µm) and type B (>0.08 µm) similarly as described in Randall *et al*. (2022). At early stages (LEP and ZYG), type A foci predominate and are significantly more abundant, with broad variance observed particularly in zygotene (Fig. S9). As meiosis progresses to pachytene, type A foci resolve into fewer, brighter type B foci, marking class I COs designation. Notably, a significant reduction in HEI10 foci is observed at pachytene across all genotypes (p = 0.001, horizontal bars) (Fig. S9). The total decrease is expected given to the coarsening hypothesis which proposes the formation of more stable clusters of HEI10 at the expense of smaller and less stable foci (Fozard *et al*., 2023; Morgan *et al*., 2021).

In pachytene the number of class I COs was significantly increased in XXY individuals compared to both *S. vulgaris* and XY *S. latifolia* (p = 0.001; Fig.S9). The moderate increase in CO number between XXXY and XXYY individuals was expected given their frequent type D (open-ring tetravalent) configuration. This likely reflects the doubled number of chromosome pairs, and potentially contribution of additional X chromosomes, paired via PAR regions allowing to have 2-3 times more COs than in XY individuals. In comparison, *S. latifolia* exhibited on average 1.65 times more HEI10 foci per cell in pachytene than *S. vulgaris* (p ≤ 0.1). The difference between *S. vulgaris* (1.09 class I CO per bivalent) or *S. latifolia* (1.24 class I CO per bivalent) XY and polyploid individuals displays a significant increase in type B foci, reaching from 1.45 to 1.69 average number of class I COs per bivalent (p = 0.001; Fig. S9).

Together, these results indicate that the number and distribution of potential COs between sex chromosomes, reflected by HEI10 foci and the structure of physical linkage, vary across pachytene configurations and are closely tied to sex chromosome dose. These patterns range from single COs in simple XY rod bivalents to multiple COs in complex multivalent configurations (Fig. 1), with implications for recombination frequency and meiotic stability.

## Discussion

In contrast to animal species, the configuration of sex chromosome during prophase I and in metaphase I has only been investigated in a few plant species. The most notable examples are in *Rumex* species, namely *Rumex acetosa, R. acetosella* (Cuñado *et al*., 2007), *R. hastatulus* (Kasjaniuk *et al*., 2019), and minor extent in *S. latifolia* (Zluvova *et al*., 2007; Warmke and Blakeslee, 1939). To address this gap, we conducted a comprehensive cytogenetic analysis of sex chromosome pairing during meiosis in *Silene*, using the advantage of recently developed tools and 3D super-resolution microscopy.

### Implications of limited sex chromosome association in higher-ploidy level

The observed architecture of meiotic chromosome pairing in *Silene* was shaped by sex chromosome composition and ploidy level. In diploid *S. vulgaris* and *S. latifolia* canonical bivalent configuration was observed, whilst the XXY, XXXY, and XXYY karyotypes displayed increasing variability in trivalent and tetravalent formation (Fig. 1). In *S. latifolia*, the consistent formation of rod bivalents between X and Y chromosomes supports previous findings of distant pairing in wild-type male *S. latifolia* (Zluvova *et al*., 2007). In this study, we clearly identified the distal pairing regions using the XY-PAR oligo-painting probe, corroborating physical association and limited recombination restricted to the PAR (Bergero *et al*., 2013; Campos *et al*., 2016; Bergero *et al*., 2008; Akagi *et al*., 2025).

In XXY individuals, the Y chromosome was typically associated with one X via the PAR, while the second X remained unpaired, forming either a univalent or, possibly, an early-separated component of a transient trivalent (Fig. 1, type B). While the latter scenario cannot be entirely excluded, the univalent interpretation is more likely given the physical separation and more frequent occurrence of type A configuration (Fig. 1, S4). This is further supported by the total number of HEI10 foci which is comparable to tetraploid plants (Fig. S9). The number of HEI10 foci suggest large number of bi-trivalent configurations, leading to potential interlocks that need to be resolved prior to completing meiosis. The higher number of interlocks in turn may impair or delay synapsis, SC continuity and chromosome structure, influencing HEI10 diffusion and coarsening (Olaya *et al*., 2024; Zickler and Kleckner, 2023; Capilla-Pérez *et al*., 2021).

Early cytogenetic studies in *Silene latifolia* reported chain quadrivalents as the most prevalent type of metaphase I association(Warmke and Blakeslee, 1939), though Westergaard (1938) did not observe any such configurations. In our study, the behavior of the Y chromosome in XXXY and XXYY individuals was in fact much more variable and configuration dependent. In open-ring tetravalents (e.g., types D and E), the Y was stably integrated into a multivalent structure through the PAR, while in other configurations (e.g., type J), it appeared only loosely associated or partially excluded from the main chromosomal complex. In chain-like tetravalents (Fig. 1, type G), the Y chromosome often occupied a terminal position, suggesting limited CO engagement and weaker integration into the SC. Thus, while our findings are consistent with those of Warmke and Blakeslee (1939) we observed a higher frequency of open-ring tetravalent configurations, possibly due to the greater number of cells scored in our analysis and increased resolution. Similarly to our results, two configurations of sex chromosomes in XYY individuals were described in *R. acetosa* (Cuñado *et al*., 2007). In the case Y-X-Y association, the two Ys pair with the X chromosome, leading to two asynapsed Y axis, while in the Y-Y-X configuration, both Ys undergo at least partial synapsis, leaving substantial part of the X chromosome asynapsed. The latter configuration in *R. acetosa* is similar to what we observed in XXYY individuals (Fig. 1, type F), though the ratio of cells with Y-X-Y and Y-Y-X configurations is currently not known. Compared to *R. acetosa* or the species with multivalent sex chromosome configurations, such as *Leptodactylus pentadactylus* (Noronha *et al*., 2020), our observations indicate that the Y chromosome is capable of synapsis through the PAR, but its pairing capacity remains strongly constrained. This became more obvious in the polyploid context with additional and potential homologous partner (Fig. 1). Therefore, we conclude that the pairing capacity, is already repressed, reflecting (i) sequence collinearity, (ii) the evolutionary history and (iii) epigenetic differences between PAR, the X chromosome and the non-recombining region of the Y (Akagi *et al*., 2025; Moraga *et al*., 2025; Bačovský *et al*., 2019). This is also supported by the evidence of uniform quadrivalent configuration in *Humulus lupulus* var. *cordifolius* that exhibits exclusively a chain-type multivalent at metaphase I, indicating a conserved pairing behavior likely driven by fixed structural rearrangements (Ono and Suzuki, 1962).

Interestingly, in *S. vulgaris* a single rod bivalent alongside two ring bivalents may represent the autosomal subgenome with pre-existing recombination patterns that will predispose it to a sex chromosome-like configuration. Indeed, comparison of *S. vulgari*s and *S. latifolia* genomes has identified four orthologous chromosomes containing regions with limited recombination (Akagi et al., 2025; Moraga et al., 2025). Therefore, we anticipate that the same chromosomes contribute to the asynaptic behaviour observed in our study (Fig. 3c–f). Phillips *et al*. (2012) described the termination of partially synapsed bivalents near the nucleolus in *H. vulgare*, indicating delayed synapsis in regions associated with nucleolar organizing regions (NORs). Although, a similar mechanism is likely at play in *S. vulgaris*, this species possesses a substantially higher number of rDNA loci (seven chromosomal pairs for 25S rDNA and two pairs for 5S rDNA) (Siroký *et al*., 2001), which could result in late synapsis or incomplete pairing during pachytene (Phillips *et al*., 2012). Therefore, the observed asynaptic behaviour of specific autosomal chromosome regions in *S. vulgaris* more closely resembles the pairing dynamics of low-recombination domains and sex chromosomes, suggesting shared structural or epigenetic constraints that impede full synapsis.

In line with the physical positioning and immunolocalization results, volumetric quantification of ASY1- and ZYP1-labeled axes across leptotene, zygotene, and pachytene stages revealed a consistent decrease in ASY1 signal and a corresponding increase in ZYP1 as synapsis progressed (Fig. 5a). This transition was observed in both *S. vulgaris* and *S. latifolia*, as well as in polyploid individuals (XXY, XXXY, and XXYY). Despite the differences in sex chromosome number and genome size, the overall trend in ASY1/ZYP1 ratio dynamics was conserved. Consistently, quantification of residual ASY1 signal across ploidy levels confirmed that additional sex chromosomes do not facilitate extended synapsis beyond the PAR, highlighting the persistent structural and epigenetic constraints governing XY interactions (Fig. 5b). The observation of large HEI10 foci during pachytene corresponds to the phenomenon observed in other species, supporting coarsening hypothesis (Morgan *et al*., 2021; Fozard *et al*., 2023). In our study, the observed average number of class I COs per bivalent as measured by HEI10 foci at pachytene, was 1.09 in *S. vulgaris* and 1.24 in *S. latifoli*a, and 1.45 to 1.69 in higher ploidy individuals. This is comparable to, or even exceeds, CO frequencies reported in reptiles such as *Sceloporus* lizards, where total COs (including both class I and II) range between 1.14 and 1.45 per bivalent in males and females (Marín-Gual *et al*., 2022). The authors concluded that the rate of COs in reptiles is low compared to that in mammals. Given that our estimates are based solely on class I COs, the actual average number of COs is likely underestimated, indicating that *Silene* does not exhibit a particularly low recombination rate per bivalent (Fig. S9). In fact, the CO ratio in male *Silene* meiosis is comparable to that observed in *A. thaliana* (Li *et al*., 2021) or maize (Luo *et al*., 2019), aligning with the typical average of 1 to 3 COs per chromosome pair found across most eukaryotic species (Brazier *et al*., 2025).

### Asynapsis during prophase I is shaped by Y chromosome architecture and epigenetic features

The Y chromosome in *S. latifolia* has undergone at least two inversion events that disrupted collinearity with the X chromosome, accelerating recombination suppression and reducing the extent of homologous pairing (Marais *et al*., 2025; Hobza *et al*., 2007; Bergero *et al*., 2008; Bergero *et al*., 2013). In addition, the large non-recombining region of the Y chromosome is depleted of active histone marks, notably H3K4me3 and H3K27me3 (Bačovský *et al*., 2019), and exhibits extensive transposable element accumulation and distinct patterns of DNA methylation (Akagi *et al*., 2025). In maize, recombination hotspots are positively correlated with histone occupancy and enrichment of H3K4me3 and H3K27me3, particularly in gene-rich euchromatic regions (Kianian *et al*., 2018; Chowdary *et al*., 2023). In *S. latifolia*, both histone modifications are substantially enriched only within the PAR of the Y chromosome and at the distal ends of the X chromosome (Bačovský *et al*., 2019). Therefore, we anticipate that both sequence homology and epigenetic compatibility are essential prerequisites for successful synapsis of sex chromosomes in *S. latifolia*, potentially explaining the observed metaphase I configurations and pachytene pairing dynamics (Fig. 5a, b).

The limited recombination near the X chromosome centromere and across the Y-specific region likely reflects not only structural divergence (Akagi *et al*., 2025) but also progressive epigenetic erosion, which may further contribute to gene down-regulation and functional decay (Bachtrog *et al*., 2011; Zhou and Bachtrog, 2012; Muyle *et al*., 2021). Thus, the epigenetic and chromatin features may shape the CO distribution and frequency and have a large impact on the non-recombining region that is anticipated. Whether the asynaptic regions identified in this study, similar to those seen in the Y–X–Y configuration of *R. acetosa*, undergo meiotic silencing of unsynapsed chromatin, a process analogous to meiotic sex chromosome inactivation (MSCI) described in animals (Turner, 2007; Alavattam *et al*., 2021), remains an open question for future investigation.

## Author contribution

VB and DP planned and designed research. VB, PN, LH and JK performed experiments. Rč and VB analysed data. JC analysed the genome size. BJ analysed the sequence data. VB wrote the main text of the manuscript with the input of all authors. All authors approved the final version of the manuscript.

## Acknowledgement

The work was supported from the project TowArds Next GENeration Crops, reg. no. CZ.02.01.01/00/22_008/0004581 of the ERDF Programme Johannes Amos Comenius. We acknowledge the core facility CELLIM supported by the Czech-BioImaging large RI project (LM2023050 funded by MEYS CR) for their support with obtaining scientific data presented in this paper.

## Supplementary information

**Supplementary Note S1**.

**Table S1**. Polyploid induction and genome size measurements.

**Table S2**. Sequence similarity overview.

**Table S3**. Number of analysed cells per each phase.

**Figure S1**. Genome size measurement of individual *Silene* plants. The genome sizes of individual plants are detailed in Tables S1.

**Figure S2**. Isolation and characterization of the *Silene* ZYP1 gene. (a) Multiple sequence alignment of ZYP1 from various *Silene* species. (b) Multiple alignment of amplified clones of the full-length ZYP1 sequence. (c) Hydrophobicity analysis of the ZYP1 protein using the Kyte & Doolittle algorithm with a linear weight variation model. (d) Predicted structure of the ZYP1 protein. (e) 3D visualization of the ZYP1 protein structure generated using AlphaFold2.

**Figure S3**. Organization of sex chromosome and autosomes at metaphase I in *S. vulgaris* and *S. latifolia*. Two satellite markers, X43.1 (subtelomeric, green) and STAR-C (centromeric, magenta), were used to distinguish sex chromosomes–specifically the Y q-arm, which bears the X43.1 satellite near the PAR. The XY-PAR oligo-painting probe (red), previously described in Bačovský et al. (2020), specifically labels the X and Y chromosomes, hybridizing to both ends of the X chromosome and within the PAR of the Y chromosome. In *S. vulgaris*, the same XY-PAR oligo painting probe identifies three orthologous chromosomes. Chromosomes were counterstained with DAPI. Scale bar = 10 μm.

**Figure S4**. Organization of sex chromosomes and autosomes at the metaphase I in *Silene latifolia* individuals with an XXY constitution (2n = 3x = 36). (a) A bivalent and univalent structure, with single X chromosome and a single Y chromosome forming a bivalent (XY-PAR positive regions, red). The second X chromosome is being physically separated forming potential self-ring univalent. (b) A Y-shaped trivalent, in which both X chromosomes are linked to the Y via the pseudoautosomal region (PAR). Two satellite markers, X43.1 (subtelomeric, green) and STAR-C (centromeric, magenta), were used to distinguish sex chromosomes– specifically the q-arm of the Y chromosome, which bears the X43.1 satellite near the PAR. The XY-PAR oligo-painting probe, previously described in Bačovský et al. (2020), specifically labels the X and Y chromosomes, hybridizing to both ends of the X chromosome and within the PAR of the Y chromosome. Chromosomes were counterstained with DAPI. Scale bar = 10 μm.

**Figure S5**. Organization of sex chromosomes and autosomes at metaphase Iin *Silene latifolia* individuals with an XXXY constitution (2n = 4x = 48). (a) A chain-like tetravalent is formed by three X chromosomes and a single Y chromosome (XY-PAR positive regions, red). (b) An open-ring tetravalent structure, resulting from two bivalent-like X chromosomes connected at both arms, with the remaining X and Y chromosomes attached via the pseudoautosomal region (XY-PAR). Two satellite markers, X43.1 (subtelomeric, green) and STAR-C (centromeric, magenta), were used to distinguish sex chromosomes–specifically q-arm of the Y chromosome, which bears the X43.1 satellite near the PAR. The XY-PAR oligo-painting probe, previously described in Bačovský et al. (2020), specifically targets X and Y chromosomes. It hybridizes to both ends of the X chromosome and within the PAR region of the Y chromosome. Chromosomes were counterstained with DAPI. Scale bar = 10 μm.

**Figure S6**. Organization of sex chromosomes and autosomes at metaphase I in *Silene latifolia* individuals with an XXYY constitution (2n = 4x = 48, XXYY). (a) An open-ring tetravalent structure formed by two X chromosomes connected at both arms, with an additional X and a Y chromosome linked via the pseudoautosomal region (XY-PAR, red). (b) A single bivalent and rod bivalent structures, with two X chromosomes paired via both arms and two Y chromosomes associated only through the PAR. (c) A chain-like tetravalent configuration, with two X chromosomes centrally positioned and two Y chromosomes connected via PAR. (d) An open-ring tetravalent in which two X chromosomes are connected through the q-arm, and two Y chromosomes are linked via PAR to one of the X chromosomes. (e) An open-ring tetravalent composed of one X and two Y chromosomes connected via the PAR, keeping one X chromosome as the main pairing partner. (f) A chain tetravalent structure with a loosely associated Y chromosome. Two satellite markers, X43.1 (subtelomeric, green) and STAR-C (centromeric, magenta), were used to distinguish sex chromosomes–particularly the Y q-arm, which bears the X43.1 signal near the PAR. The XY-PAR oligo-painting probe (red), previously described in Bačovský et al. (2020), specifically labels the X and Y chromosomes, hybridizing to both ends of the X chromosome and within the PAR region of the Y chromosome. Chromosomes were counterstained with DAPI. Scale bar = 10 μm.

**Figure S7**. Progression of meiosis from leptotene to pachytene in triploid male *Silene latifolia* plants (2n = 3x = 36) with XXY sex chromosome constitution. The localization and dynamics of ZYP1, ASY1, and HEI10 were consistent across all analysed cells (n = 26; Table S3), based on six independent experiments. Scale bar = 4 μm.

**Figure S8**. Progression of meiosis from leptotene to pachytene in male autotetraploid *Silene latifolia* individuals. Panels (a, c, e) show individuals with an XXYY sex chromosome constitution, while panels (b, d, f) depict individuals with an XXXY constitution. The localization patterns of ZYP1, ASY1, and HEI10 were consistent across all analysed cells (n = 52; Table S3) from at least six independent experiments. Scale bar = 4 μm.

**Figure S9**. Quantification of HEI10 foci classified as type A (early, diffuse foci, blue) and type B (late, large foci, yellow) across leptotene, zygotene and pachytene. HEI10 foci were classified in *S. vulgaris* and *S. latifolia* with different sex chromosome constitutions: diploid (2n = 2x = 24, XY), triploid (2n = 3x = 36, XXY), and autotetraploid (2n = 4x = 48, XXXY or XXYY). Each dot represents the number of foci counted per cell (n per group indicated in Table S3). The CO were counted in cells with non-disrupted ZYP1 elements and mature HEI10 foci. *** p ≤ 0.001, ** p ≤ 0.01, * p ≤ 0.05, · p ≤ 0.1 values were obtained by one-way ANOVA followed by Tukey’s multiple comparisons.

