## Supplementary information for "Sex chromosome pairing and multivalent associations during meiosis in diploid and polyploid *Silene latifolia*"

**List of items:**

Supplementary Note S1

**Table S1.** Polyploid induction and genome size measurements.

**Table S2.** Sequence similarity overview.

**Table S3.** Number of analysed cells per each phase.

**Figure S1.** Genome size measurement of individual *Silene* plants.

**Figure S2.** Isolation and characterization of the *Silene* ZYP1 gene.

**Figure S3.** Organization of sex chromosome and autosomes at metaphase I in *S. vulgaris* and *S. latifolia*.

**Figure S4.** Organization of sex chromosomes and autosomes at the metaphase I in *Silene latifolia* individuals with an XXY constitution (2n = 3x = 36).

**Figure S5.** Organization of sex chromosomes and autosomes at metaphase Iin *Silene latifolia* individuals with an XXXY constitution (2n = 4x = 48).

**Figure S6.** Organization of sex chromosomes and autosomes at metaphase I in *Silene latifolia* individuals with an XXYY constitution (2n = 4x = 48, XXYY).

**Figure S7.** Progression of meiosis from leptotene to pachytene in triploid male *Silene latifolia* plants (2n = 3x = 36) with XXY sex chromosome constitution.

**Figure S8.** Progression of meiosis from leptotene to pachytene in male autotetraploid *Silene latifolia* individuals.

**Figure S9.** Quantification of HEI10 foci classified as type A (early, diffuse foci, blue) and type B (late, large foci, yellow) across leptotene, zygotene and pachytene.

**Supplementary Note S1**

Seedlings at the cotyledon stage were treated at 3-day intervals or left untreated as a control. A single drop of warm colchicine or oryzalin solution was pipetted onto the cotyledons of each seedling (52 plants per treatment) to cover the emerging shoot. The seedlings were then placed in a high humidity environment (approximately 80-90% of relative humidity). After the treatment, plants were grown under standard conditions for 14 days. Due to the low survival rate (1.92-3.85%) of treated plants compared to control (96%), we applied 5 mM colchicine to adult female flowers 2 days after pollination (2DAP). As a control for this treatment, we applied deionized water without colchicine. From 36 tested individuals, 6 polyploid plants were successfully generated and further analysed under the light microscope. Plants possessing 2n=4x=48, XXYY were selected for generating the F2 generation, which was analysed again by flow cytometry and chromosome squashing.

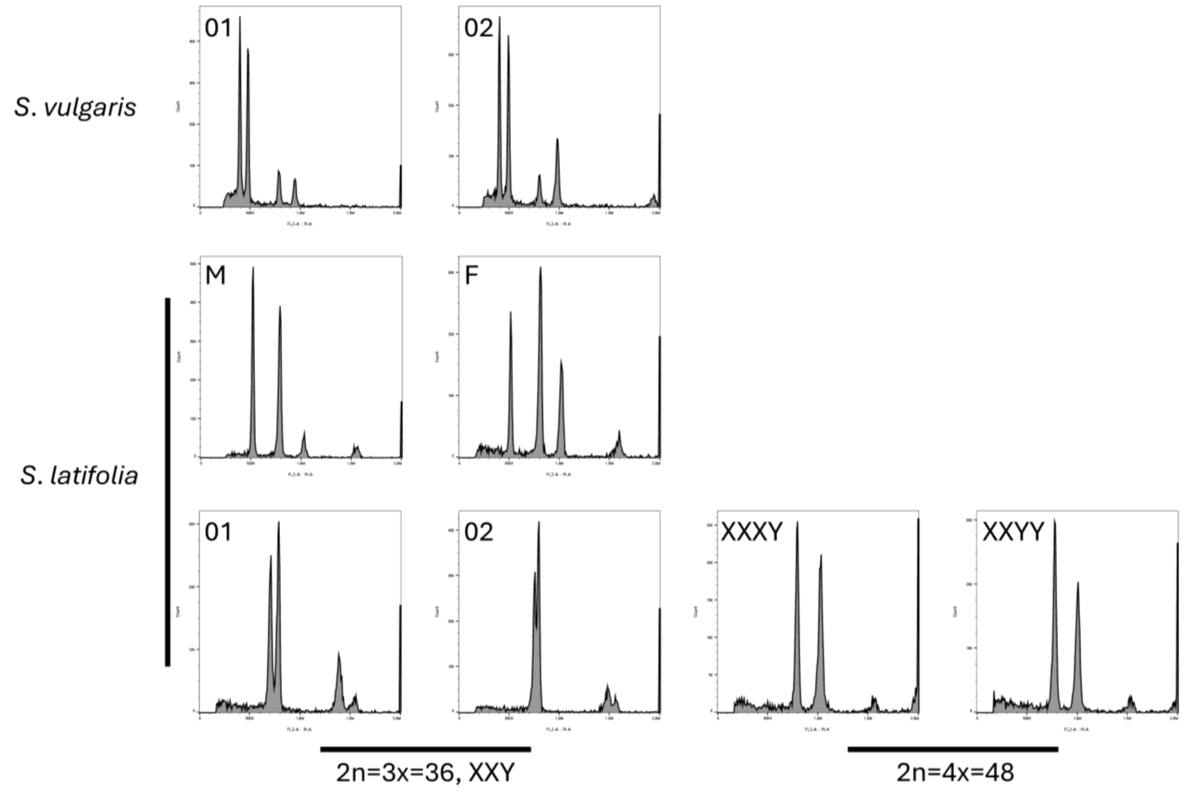


**Figure S1. Genome size measurement of individual *Silene* plants.** The genome sizes of individual plants are detailed in Tables S1.

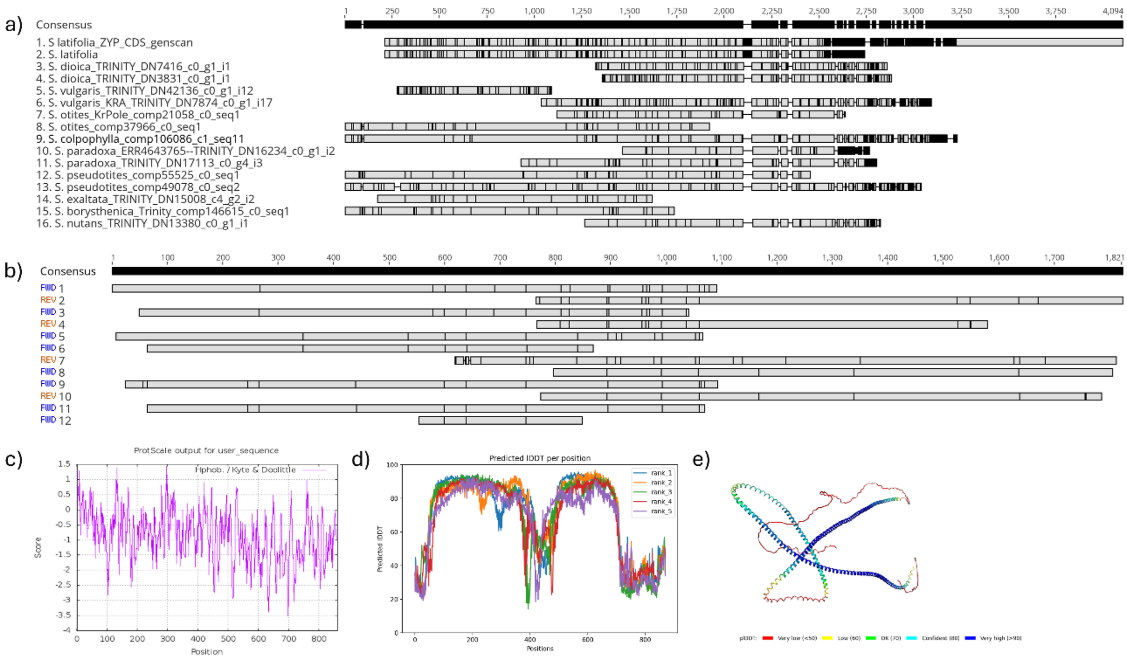


**Figure S2. Isolation and characterization of the *Silene* ZYP1 gene.** (a) Multiple sequence alignment of ZYP1 from various *Silene* species. (b) Multiple alignment of amplified clones of the full-length ZYP1 sequence. (c) Hydrophobicity analysis of the ZYP1 protein using the Kyte & Doolittle algorithm with a linear weight variation model. (d) Predicted structure of the ZYP1 protein. (e) 3D visualization of the ZYP1 protein structure generated using AlphaFold2.

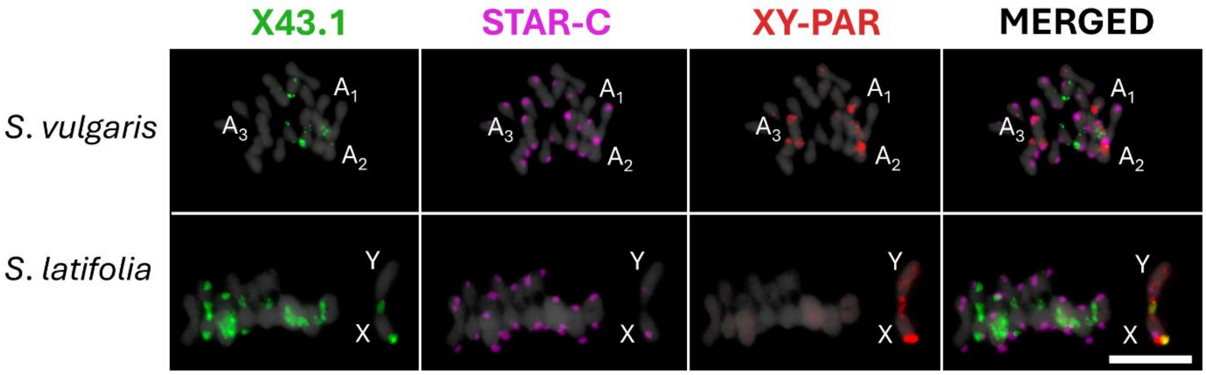


**Figure S3.** Organization of sex chromosome and autosomes at metaphase I in *S. vulgaris* and *S. latifolia*. Two satellite markers, X43.1 (subtelomeric, green) and STAR-C (centromeric, magenta), were used to distinguish sex chromosomes–specifically the Y q-arm, which bears the X43.1 satellite near the PAR. The XY-PAR oligo-painting probe (red), previously described in Bačovský et al. (2020), specifically labels the X and Y chromosomes, hybridizing to both ends of the X chromosome and within the PAR of the Y chromosome. In *S. vulgaris*, the same XY-PAR oligo painting probe identifies three orthologous chromosomes. Chromosomes were counterstained with DAPI. Scale bar = 10 µm.

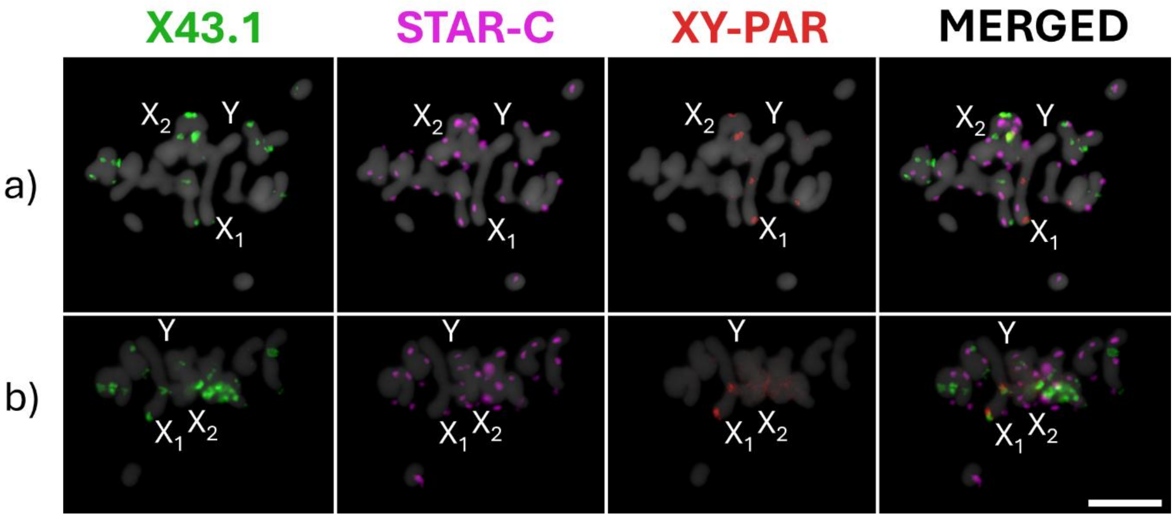


**Figure S4.** Organization of sex chromosomes and autosomes at the metaphase I in *Silene latifolia* individuals with an XXY constitution (2n = 3x = 36). (a) A bivalent and univalent structure, with single X chromosome and a single Y chromosome forming a bivalent (XY-PAR positive regions, red). The second X chromosome is being physically separated forming potential self-ring univalent. (b) A Y-shaped trivalent, in which both X chromosomes are linked to the Y via the pseudoautosomal region (PAR). Two satellite markers, X43.1 (subtelomeric, green) and STAR-C (centromeric, magenta), were used to distinguish sex chromosomes–specifically the q-arm of the Y chromosome, which bears the X43.1 satellite near the PAR. The XY-PAR oligo-painting probe, previously described in Bačovský et al. (2020), specifically labels the X and Y chromosomes, hybridizing to both ends of the X chromosome and within the PAR of the Y chromosome. Chromosomes were counterstained with DAPI. Scale bar = 10 µm.

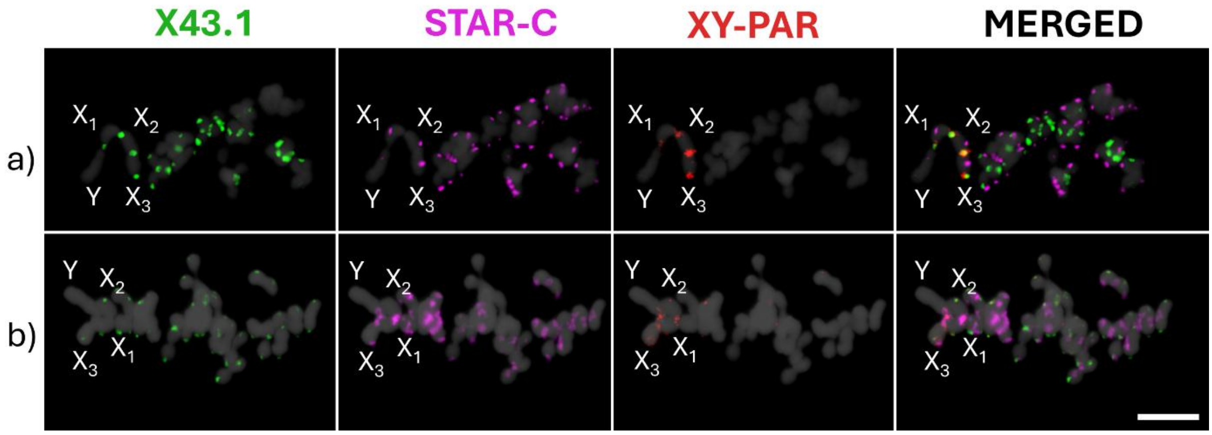


**Figure S5.** Organization of sex chromosomes and autosomes at metaphase Iin *Silene latifolia* individuals with an XXXY constitution (2n = 4x = 48). (a) A chain-like tetravalent is formed by three X chromosomes and a single Y chromosome (XY-PAR positive regions, red). (b) An open-ring tetravalent structure, resulting from two bivalent-like X chromosomes connected at both arms, with the remaining X and Y chromosomes attached via the pseudoautosomal region (XY-PAR). Two satellite markers,

X43.1 (subtelomeric, green) and STAR-C (centromeric, magenta), were used to distinguish sex chromosomes–specifically q-arm of the Y chromosome, which bears the X43.1 satellite near the PAR. The XY-PAR oligo-painting probe, previously described in Bačovský et al. (2020), specifically targets X and Y chromosomes. It hybridizes to both ends of the X chromosome and within the PAR region of the Y chromosome. Chromosomes were counterstained with DAPI. Scale bar = 10 µm.

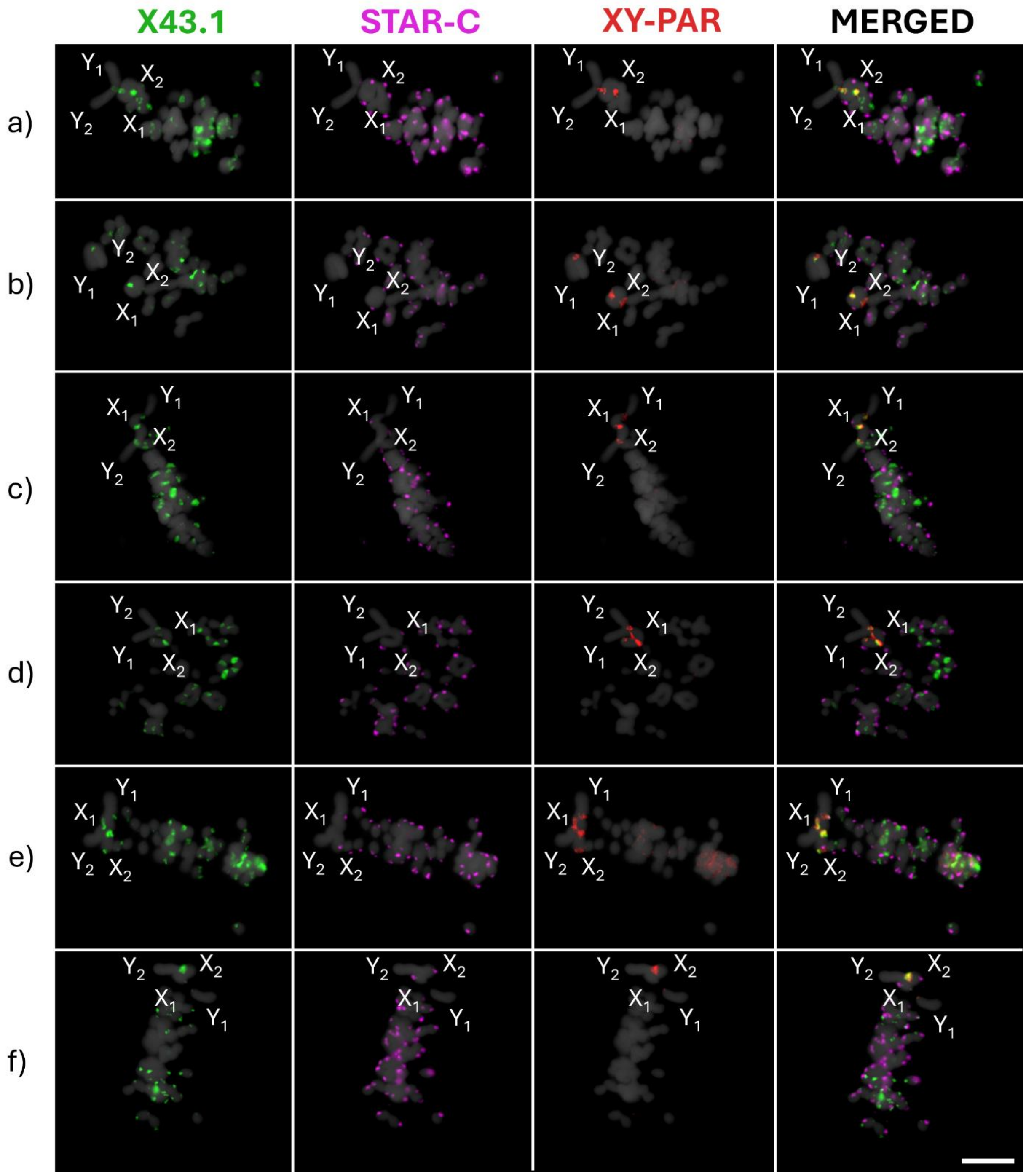


**Figure S6.** Organization of sex chromosomes and autosomes at metaphase I in *Silene latifolia* individuals with an XXYY constitution (2n = 4x = 48, XXYY). (a) An open-ring tetravalent structure formed by two X chromosomes connected at both arms, with an additional X and a Y chromosome linked via the pseudoautosomal region (XY-PAR, red). (b) A single bivalent and rod bivalent structures, with two X chromosomes paired via both arms and two Y chromosomes associated only through the PAR. (c) A chain-like tetravalent configuration, with two X chromosomes centrally positioned and two Y chromosomes connected via PAR. (d) An open-ring tetravalent in which two X chromosomes are connected through the q-arm, and two Y chromosomes are linked via PAR to one of the X chromosomes. (e) An open-ring tetravalent composed of one X and two Y chromosomes connected via the PAR, keeping one X chromosome as the main pairing partner. (f) A chain tetravalent structure with a loosely associated Y chromosome. Two satellite markers, X43.1 (subtelomeric, green) and STAR-C (centromeric, magenta), were used to distinguish sex chromosomes–particularly the Y q-arm, which bears the X43.1 signal near the PAR. The XY-PAR oligo-painting probe (red), previously described in Bačovský et al. (2020), specifically labels the X and Y chromosomes, hybridizing to both ends of the X chromosome and within the PAR region of the Y chromosome. Chromosomes were counterstained with DAPI. Scale bar = 10 µm.

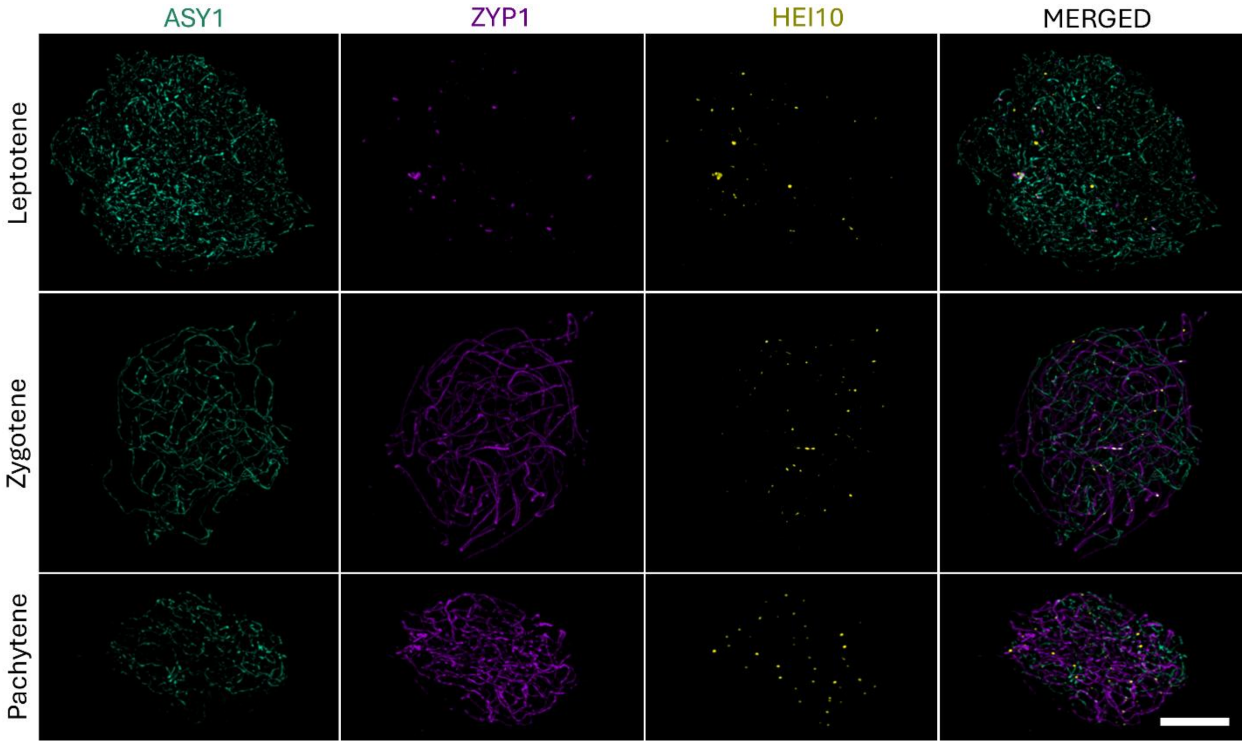


**Figure S7.** Progression of meiosis from leptotene to pachytene in triploid male *Silene latifolia* plants (2n = 3x = 36) with XXY sex chromosome constitution. The localization and dynamics of ZYP1, ASY1, and HEI10 were consistent across all analysed cells (n = 26; Table S3), based on six independent experiments. Scale bar = 4 µm.

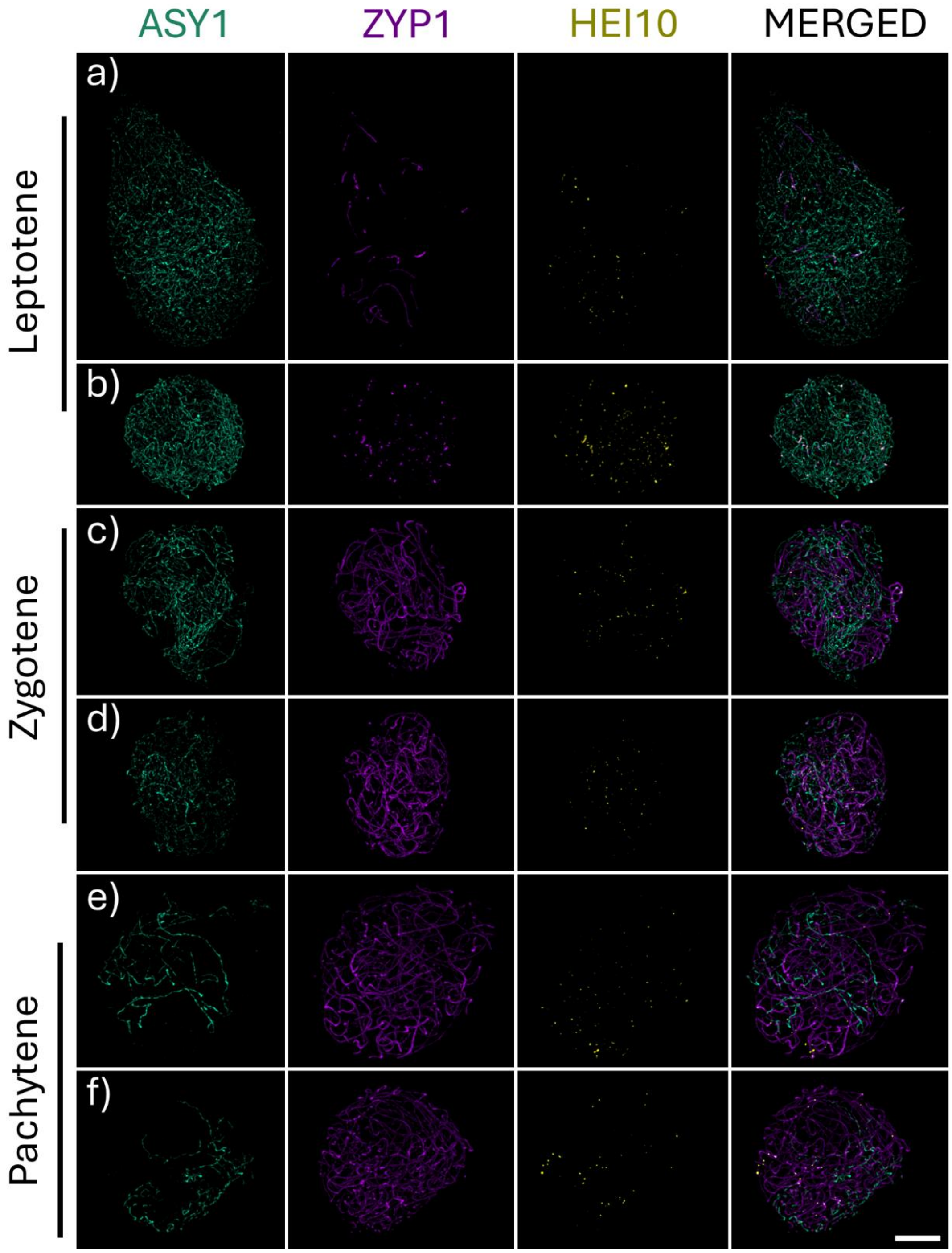


**Figure S8.** Progression of meiosis from leptotene to pachytene in male autotetraploid *Silene latifolia* individuals. Panels (a, c, e) show individuals with an XXYY sex chromosome constitution, while panels (b, d, f) depict individuals with an XXXY constitution. The localization patterns of ZYP1, ASY1, and HEI10 were consistent across all analysed cells (n = 52; Table S3) from at least six independent experiments. Scale bar = 4 µm.

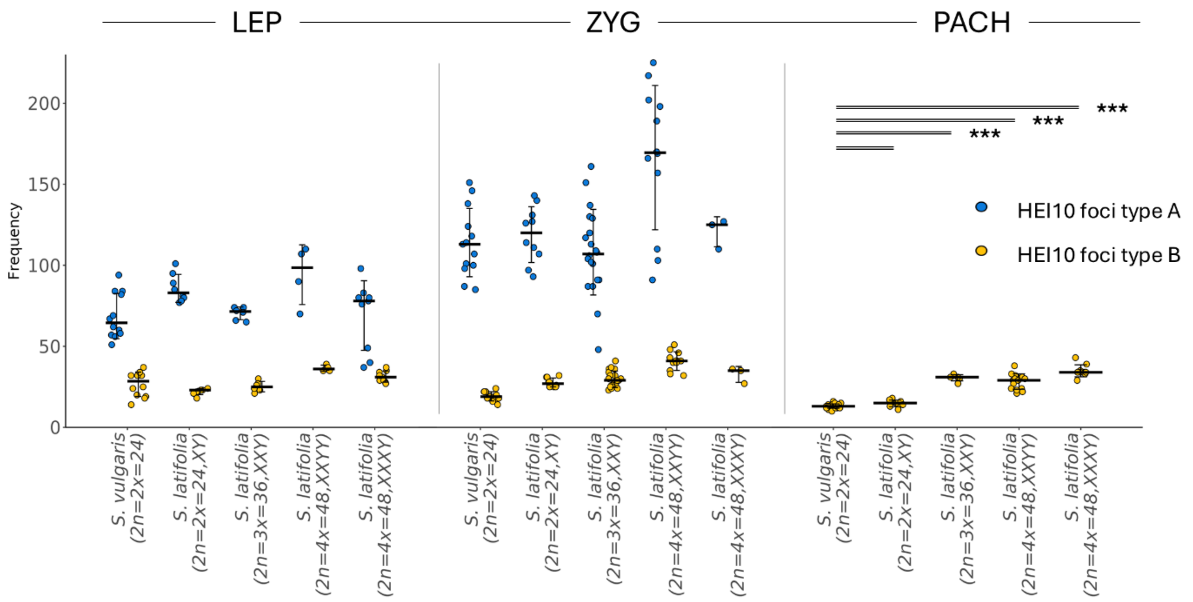


**Figure S9.** Quantification of HEI10 foci classified as type A (early, diffuse foci, blue) and type B (late, large foci, yellow) across leptotene, zygotene and pachytene. HEI10 foci were classified in *S. vulgaris* and *S. latifolia* with different sex chromosome constitutions: diploid (2n = 2x = 24, XY), triploid (2n = 3x = 36, XXY), and autotetraploid (2n = 4x = 48, XXXY or XXYY). Each dot represents the number of foci counted per cell (n per group indicated in Table S3). The CO were counted in cells with non-disrupted ZYP1 elements and mature HEI10 foci. *** $p \leq 0.001$, ** $p \leq 0.01$, * $p \leq 0.05$, · $p \leq 0.1$ values were obtained by one-way ANOVA followed by Tukey's multiple comparisons.

Table S3. Number of analysed cells per each phase.

| Plant | Ploidy level | | Leptotene | Zygotene | Pachytene |
|---|---|---|---|---|---|
| S. vulgaris | 2n=2x=24 | Number of cells analysed | 10 | 13 | 24 |
| S. latifolia | 2n=2x=24, XY | | 8 | 10 | 9 |
| | 2n=3x=36, XXY | | 6 | 20 | 7 |
| | 2n=4x=48, XXYY | | 5 | 12 | 15 |
| | 2n=4x=48, XXXY | | 7 | 3 | 10 |
